# A Hybrid *de novo* Assembly of the Sea Pansy (*Renilla muelleri*) Genome

**DOI:** 10.1101/424614

**Authors:** Justin Jiang, Andrea M. Quattrini, Warren R. Francis, Joseph F. Ryan, Estefanía Rodríguez, Catherine S. McFadden

## Abstract

**Background:** Over 3,000 species of octocorals (Cnidaria, Anthozoa) inhabit an expansive range of environments, from shallow tropical seas to the deep-ocean floor. They are important foundation species that create coral “forests” which provide unique niches and three-dimensional living space for other organisms. The octocoral genus *Renilla* inhabits sandy, continental shelves in the subtropical and tropical Atlantic and eastern Pacific Oceans. *Renilla* is especially interesting because it produces secondary metabolites for defense, exhibits bioluminescence, and produces a luciferase that is widely used in dual-reporter assays in molecular biology. Although several cnidarian genomes are currently available, the majority are from hexacorals. Here, we present a *de novo* assembly of the *R. muelleri* genome, making this the first complete draft genome from an octocoral.

**Findings:** We generated a hybrid *de novo* assembly using the Maryland Super-Read Celera Assembler v.3.2.6 (MaSuRCA). The final assembly included 4,825 scaffolds and a haploid genome size of 172 Mb. A BUSCO assessment found 88% of metazoan orthologs present in the genome. An Augustus *ab initio* gene prediction found 23,660 genes, of which 66% (15,635) had detectable similarity to annotated genes from the starlet sea anemone, *Nematostella vectensis,* or to the Uniprot database. Although the *R. muelleri* genome is smaller (172 Mb) than other publicly available, hexacoral genomes (256-448 Mb), the *R. muelleri* genome is similar to the hexacoral genomes in terms of the number of complete metazoan BUSCOs and predicted gene models.

**Conclusions:** The *R. muelleri* hybrid genome provides a novel resource for researchers to investigate the evolution of genes and gene families within Octocorallia and more widely across Anthozoa. It will be a key resource for future comparative genomics with other corals and for understanding the genomic basis of coral diversity.

## Data Description

### Organism Description

Octocorallia is a subclass of Anthozoa (Phylum: Cnidaria) that is comprised of three orders: Alcyonacea, Helioporacea, and Pennatulacea [1]. The Pennatulacea, commonly known as sea pens, are a monophyletic group [1, 2] and are the most morphologically distinct group of octocorals [1, 3]. Sea pens differ from other octocorals by exhibiting the most integrated colonial behavior, with colonies arising from an axial polyp that develops into a peduncle—used to anchor the animal into soft-sediments or onto hard surfaces—and a rachis that supports secondary polyps [1, 3-4]. There are 14 valid families of Pennatulacea distinguished by the arrangement of the secondary polyps around the rachis [1, 4]. The monogeneric family Renillidae Lamarck, 1816 consists of seven species [5], unique because of their foliate colony growth form [1, 4].

*Renilla* is found naturally on sandy, shallow sea floors along the Atlantic and Pacific coasts of North and South America [3, 4, 6]. The brilliant bioluminescence and endogenous fluorescence of these animals have led to them becoming important organisms in microscopy and molecular biology. Isolated originally from *R. reniformis*, the enzyme luciferase (Renilla-luciferin 2-monooxygenase) is used in dual luciferase reporter assays, which are commonly used to study gene regulation and expression, signaling pathways, and the structure of regulatory genes [7-8]. The green fluorescent protein from *Renilla* has medical applications as well as general molecular biology and imagery uses [9]. In addition, the compounds produced by *Renilla* for chemical defense [10] may be important sources for discovery of marine natural products [11]. Thus, a genome of the octocoral *Renilla* is highly valuable to the scientific community, providing a novel resource that has a range of important uses—from molecular biology to comparative genomics.

Due to the known difficulties of resolving lengthy repeat regions with Illumina-only data [12-13], we used a hybrid assembly approach [13-14], combining long-read Pacific Biosciences (PacBio) data with short-read Illumina data. Studies have shown that a hybrid approach results in a more complete assembly with less genome fragmentation [15-17]. Our hybrid approach used low coverage PacBio reads (15x coverage) along with high coverage Illumina HiSeq reads (105x coverage) to assemble a draft genome of *R. muelleri* Schultze in Kölliker, 1872, a sea pen common to shallow waters of the Gulf of Mexico [6].

## Methods and Results

### Data Collection

A live specimen of *R. muelleri* was obtained from Gulf Specimen Marine Lab (Panacea, FL, USA), which collects specimens off the panhandle of Florida in the Gulf of Mexico. Upon receiving the specimen, it was flash frozen in liquid nitrogen. Genomic DNA was then extracted using a modified CTAB protocol [18]. A total of 5.6 µg of DNA was sent to Novogene (Sacramento, CA, USA) for library preparation and sequencing. 350 bp insert DNA libraries that were PCR free were prepared and then multiplexed with other organisms on two lanes of an Illumina HiSeq 2500 (150 bp PE reads). In addition, Illumina MiSeq and PacBio sequencing were performed at the Weill Cornell Medicine Epigenomics Core Facility in New York. For the Illumina MiSeq run, the *Renilla* library was prepared with TruSeq LT and then multiplexed with eight other corals and sequenced (300 bp PE reads, MiSeq v3 Reagent kit). For PacBio sequencing, a DNA library was prepared from 5 ug of DNA using the SMRTbell template prep kit v 1.0. Sequencing was carried out on 10 SMRT cells on a RSII instrument using P6-C4 chemistry. PacBio SMRT Analysis 2.3 subread filtering module was used to produce the subread files for assembly.

As part of another study, we sequenced total RNA from a congeneric species, *R. reniformis*. The specimen was collected alive on the beach in North Flagler County, Florida, USA. RNA was extracted from the whole adult colony and sequenced on a NextSeq500 (150 bp PE reads) instrument. Library preparation and sequencing were performed at the University of Florida’s Interdisciplinary Center for Biotechnology.

### DNA Read Processing

A total of 246,744,426 PE reads were obtained from the HiSeq and 6,725,072 PE reads were obtained from the MiSeq. In total, we generated 39,029,185,500 bases of Illumina data. Adapters were trimmed from all raw Illumina reads using Trimmomatic v.0.35 (*ILLUMINACLIP:2:30:10 LEADING:5 TRAILING:5 SLIDINGWINDOW:4:20 MINLEN:3*; Trimmomatic, RRID:SCR_011848) [19], resulting in 38.98 Gb of reads. These reads were then filtered with Kraken v.1.0 (Kraken, RRID:SCR_005484) [20] using the MiniKraken 8GB database [21] to screen for possible microbial, viral and archaeal contamination. A total of 960 Mb were removed from the read files, resulting in 36.23 Gb of 150 bp reads and 1.79 Gb of 300 bp reads.

A total of 1,227,306 PacBio subreads were obtained and screened against the NCBI environmental nucleotide database (env_nt.00 to env_nt.23) [22] using BLASTn v.2.2.31 (-*evalue 1e-10, -out_fmt 5*, RRID:SCR_001598) [23] to identify reads with environmental contaminants. The subreads that did not contain contaminants were extracted using MEtaGenome ANalyzer v.6.4.16 (MEGAN, RRID:SCR_011942) [24-25], resulting in 5.22 Gb in 1,195,521 reads.

### RNA-Seq Read Processing

We generated 119,604,588 PE reads of RNA-Seq data. We used Trimmomatic (version 0.36 (-phred33, ILLUMINACLIP:/usr/local/Trimmomatic-0.32/adapters/TruSeq3-PE.fa:2:30:12:1:true, MINLEN:36; Trimmomatic, RRID:SCR_011848) [19] to remove Illumina adapters. Trinity v 2.4.0 (--seqType fq --max_memory 250G --CPU 6 --left trim.R1.fq --right trim.R2.fq --full_cleanup; Trinity RRID_SCR:013048) [26] was used to assemble the transcriptome.

### Hybrid Genome Assembly

Two hybrid *de novo* assemblies were performed, one with the Maryland Super-Read Celera Assembler v.3.2.6 (MaSuRCA, RRID:SCR_010691) [27] and the other with SPAdes v.3.11.0 (SPAdes, RRID:SCR_000131; *k-mer lengths 21,33,55,77*) [28]. The Benchmarking Universal Single-Copy Orthologs v.3.0.2 (BUSCO, RRID:SCR_015008) [29] program with default settings was used to screen the *Renilla* genome assemblies for 978 orthologs from the Metazoan dataset as a method to evaluate the completeness of each assembly. BUSCO used BLAST v.2.2.31 [23] and HMMER v.3.1.b2 (HMMER, RRID:SCR_005305) [30] in its pipeline. The stats.sh program from BBMAP v.36.14 (bbmap) [31] was used to generate general assembly statistics for genomes produced by both programs (Table 1).

**Table 1.**
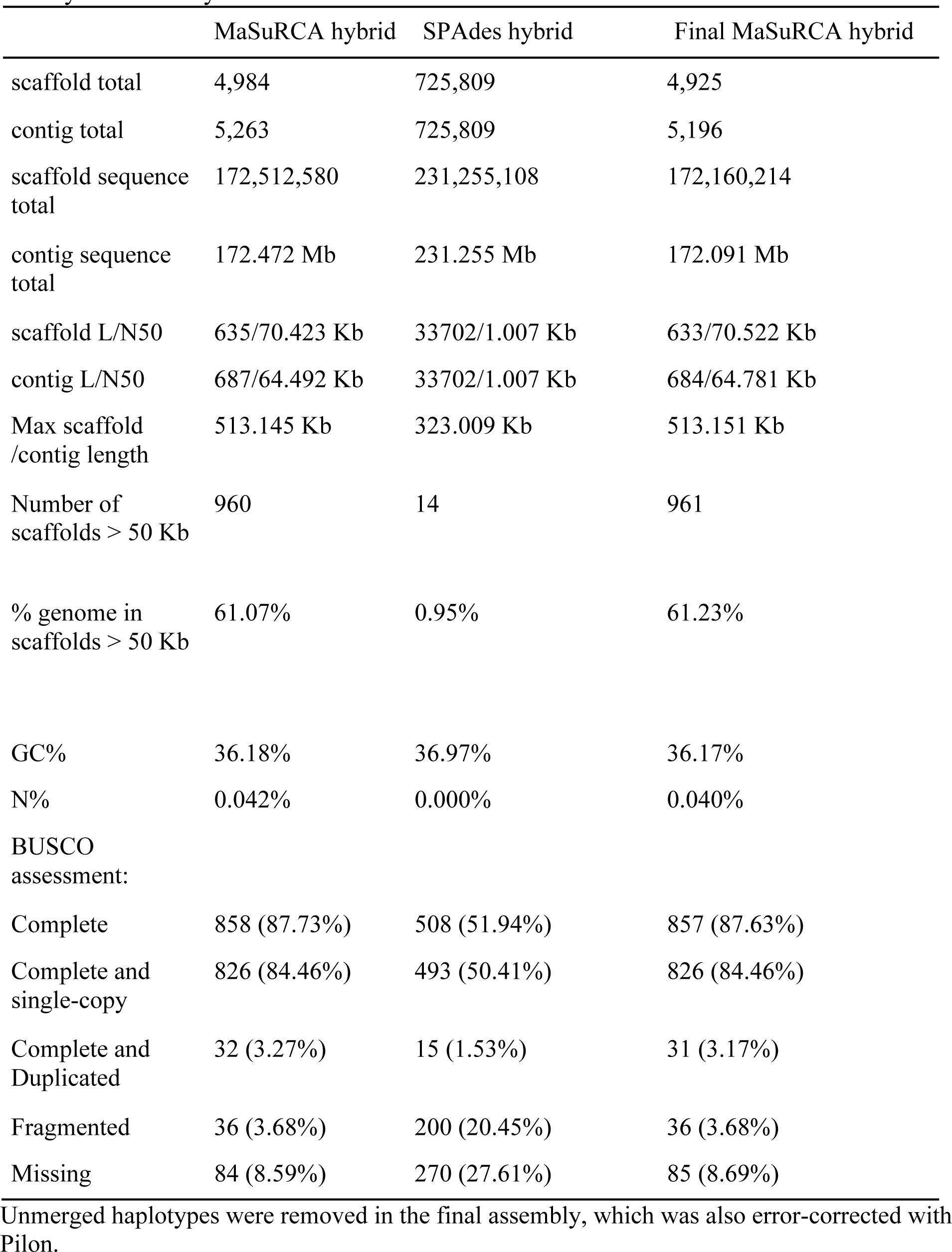
General statistics and BUSCO-completeness of both initial hybrid assemblies and the final hybrid assembly.

The MaSuRCA assembly resulted in a 147-fold decrease in the number of scaffolds generated, and a 70-fold increase in the N50 contig size (70.522 KB) as compared to the SPades assembly (1.007 KB); it also had more complete BUSCOs present (Table 1). Other statistics also indicate that the MaSuRCA assembly is much less fragmented than the SPAdes assembly (Table 1). Therefore, we used the MaSuRCA assembly in further analyses.

To improve the quality of the draft MaSuRCA assembly, six iterations of Pilon v.1.21 (Pilon, RRID:SCR_014731) [32] were used to fix assembly errors and fill assembly gaps. Bowtie2 v.2.3.2 (Bowtie2, RRID:SCR_016368) [33] was used to align Illumina HiSeq and Illumina MiSeq genomic reads to the draft assembly, and the resulting alignments were input to Pilon, which was run on default settings. A total of 52,668 SNPs were corrected, along with 14,702 small insertions and 11,841 small deletions (Supplementary Table S1).

**Supplemental Table S1.**
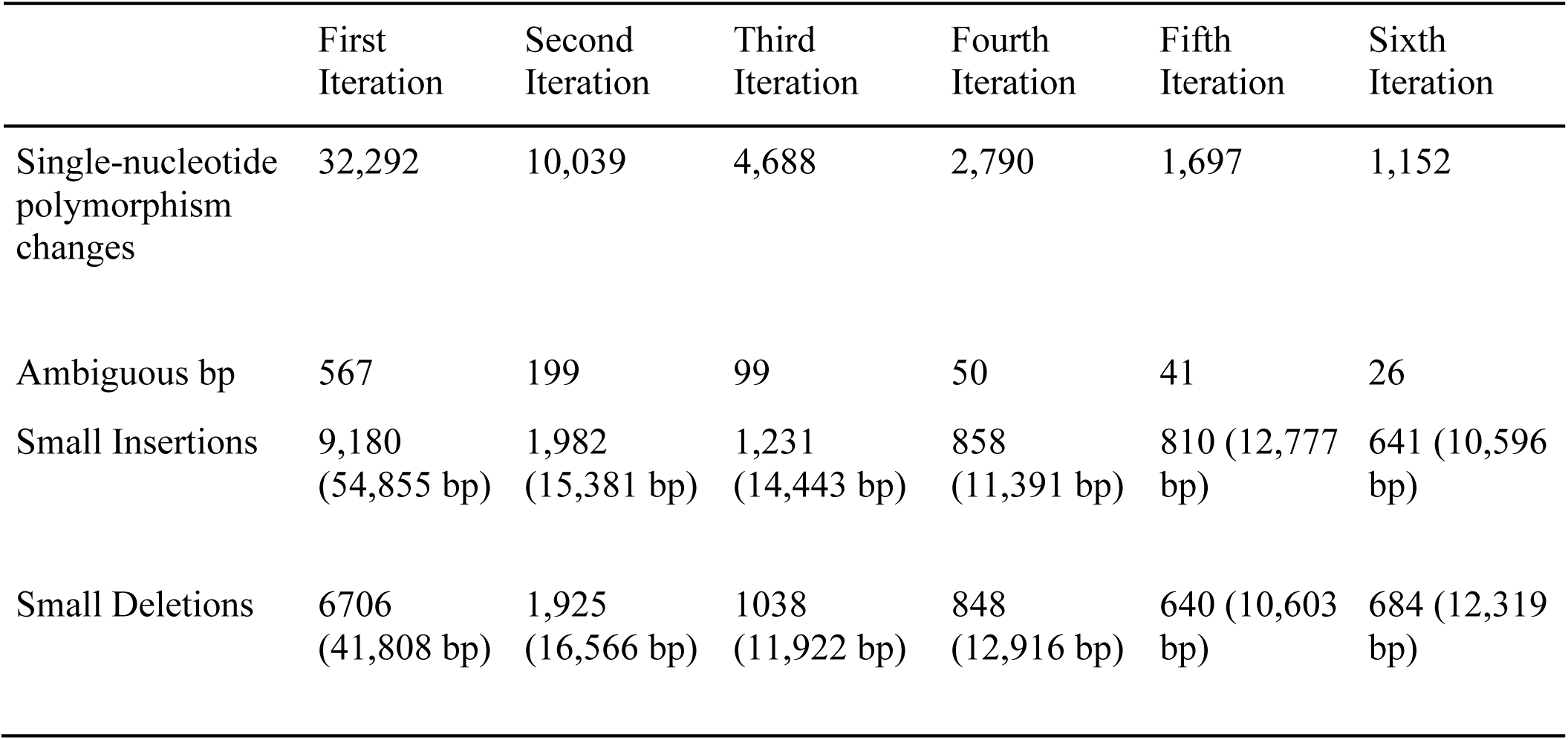
Summary of Pilon changes per iteration

To remove haploid contigs that were not merged during assembly, we ran BLASTn against the contigs themselves *(-max_target_seqs 10, -evalue 1e-40*) to find contigs that were highly similar. The custom script *haplotypeblastn.py* version 1.0 [34] filtered the BLASTn results by flagging matches that were greater than 75% identical and longer than 500 bp in length. The contigs that were identified as unmerged were subsequently removed using the *select_contigs.pl* script [35]. A total of 59 scaffolds, which amounted to 67 contigs and 384 kb, were removed from the assembly.

The bbmap program stats.sh was used to generate assembly statistics on the haplotype-removed assembly [i.e., “final assembly”, (Table 1)]. BUSCO analysis using the metazoan orthologs was again used to estimate the completeness of the final assembly, with the flag -*long* to produce higher quality training data for the downstream annotation. 857 (87.63%) orthologs were present in the final assembly (Table 1). This final *R. muelleri* assembly was masked, using RepeatMasker v.open-4.0.6 (*-species eukaryota -gccalc -div 50;* RepeatMasker, RRID:SCR_012954) [36], for downstream gene annotation. The final annotation consists of 172,512,580 bp in 4,925 scaffolds.

### Genome Annotation

Stampy v.1.0.31 (Stampy, RRID:SCR_005504) [37] was used to align 18.06 Gb of RNA-Seq data from *R. reniformis* to the masked genome to generate intron hints. The resulting bam file was processed by filtering out raw alignments using *filterBam* [38] per the recommended Augustus procedures [39]. A total of 1,837,637 intron hints were generated.

Augustus v.3.3 (*--UTR=off --allow_hinted_splicesites=atac --alternatives-from-evidence=true;* Augustus, RRID:SCR_008417) [40] was used to predict a gene model for *R. muelleri*. Augustus training was performed with the hint data from *R. reniformis*, as it has been shown to improve *ab initio* predictions [40-41]. The BUSCO-generated training data was also included to help predict a gene model. A modified extrinsic weight file was used in Augustus to penalize predicted introns that were unsupported by hint evidence and reward predicted introns that were supported by hint evidence by 1e2.

Augustus predicted 23,660 genes that had an average exon length of 249 bp and an average intron length of 524 bp as calculated by *gfstats.py* [42] (Table 2). BUSCO with the metazoan lineage (*-m prot*) orthologs was used to assess the quality of the prediction, finding 84.87% (830/978) orthologs (Table 2).

**Table 2.** Statistics for the gene model predicted by Augustus.

|  | Number |
| --- | --- |
| Genes | 23,660 |
| Exons | 140,384 |
| Introns | 117,838 |
| Average Exon Length | 249 |
| Exons Per Gene | 5.93 |
| Average Intron Length | 524 |
| Introns Per Gene | 4.98 |
| BUSCO assessment: |  |
| Complete | 830 (84.87%) |
| Complete and single-copy | 798 (81.60%) |
| Complete and Duplicated | 32 (3.27%) |
| Fragmented | 64 (6.54%) |
| Missing | 84 (8.59%) |

### Functional Annotations

BLASTp v.2.2.31+ (*-evalue 1e-10 -seg yes -soft_masking true -lcase_masking*, BLASTp, RRID:SCR_001010) [23] was used to map the predicted gene models of *R. muelleri* to filtered protein models of another anthozoan, the sea anemone, *Nematostella vectensis* (Joint Genome Institute, JGI, v 1.0) [43]. A total of 63% (14,931) of the predicted genes (23,660) mapped to *N. vectensis* proteins (27,273). A custom python script, *filterGenes.py* [44] was used to filter the matches by selecting the highest bit score; in cases where bit scores were identical, the match with the highest percent length of all matches was used as a tiebreaker. Of the 14,931 genes that mapped to *N. vectensis* proteins, 12,279 genes were annotated with GO function, KOG function and/or InterPro domains; 8,101 genes were assigned GO terms; 11,067 genes were assigned KOG functions; and 10,126 genes were assigned InterPro domains (Supplemental File 1). The 8,729 genes that did not hit *N. vectensis* proteins were remapped with BLASTp using a lower evalue (*1e-5*) and filtered with the aforementioned python script with the same settings; an additional 2,002 of the genes mapped to *N. vectensis.* Of these, 1,512 genes were annotated with GO functions, KOG functions and/or InterPro domains (Supplemental File 1). The remaining 6,727 genes that did not match *N. vectensis* annotations were mapped to the UniProt database (UniProt, RRID:SCR_002380) [45-46] with BLASTp (*-evalue 1e-5*), and 1,844 of these were assigned a UniProt function. In total, 79.36% (18,777/23,660) of the predicted gene models were mapped to either *N. vectensis* predicted proteins or the UniProt database, and 66.08% (15,635/23,660) of the predicted *Renilla* genes have either functional annotations or InterPro domain information associated with them.

We also used BLASTp (*-evalue 1e-10 -seg yes -soft_masking true -lcase_masking)* to map the predicted genes against a newer *N. vectensis* dataset that was generated using RNA-Seq (hereafter called the Vienna dataset) [47-48]. A total of 63% (15,001) of the predicted genes (23,660) mapped to the Vienna dataset (25,729) (Supplemental File 2). As above, the predicted genes that did not map were remapped with a lower e-value (*1e-5),* resulting in 2,071 additional predicted genes mapping to the *N. vectensis* Vienna dataset. In total, 72.15% (17,072) of predicted genes mapped to the Vienna dataset. This dataset did not have associated functional annotations. Combining all gene model annotation methods, 79.82% (18,886) of genes from the Augustus gene model were mapped to the JGI *N. vectensis* annotations, the *N. vectensis* Vienna dataset, or the UniProt database (Supplemental Files 1-3).

### Genome Assembly Comparisons

We compared the *R. muelleri* genome assembly to previously published anthozoan (e.g., corals, anemones) genomes using a variety of assessment statistics (Supplemental Table S2). BUSCO was used (*-m geno)* to assess the completeness of six hexacoral genomes and a draft *R. reniformis* genome and compare these results to the hybrid *R. muelleri* assembly (Fig. 1). We found the BUSCO-completeness of our *R. muelleri* assembly (857) to be more similar to the well-curated assembly of the model organism *N. vectensis* (893) [49-50] than to the other anthozoans. BUSCOs from the other five hexacoral genomes were less complete, with complete BUSCOs ranging from 728 (*Acropora digitifera*) to 839 (*Discosoma* sp.) [50-57]. Only 800 complete BUSCOs were recovered from the other hybrid assembly, the hexacoral *Montastraea cavernosa* [57]. The only other publicly-available octocoral genome, *R. reniformis*, had considerably fewer complete BUSCOs (356, Fig.1) [58].

**Figure 1.**
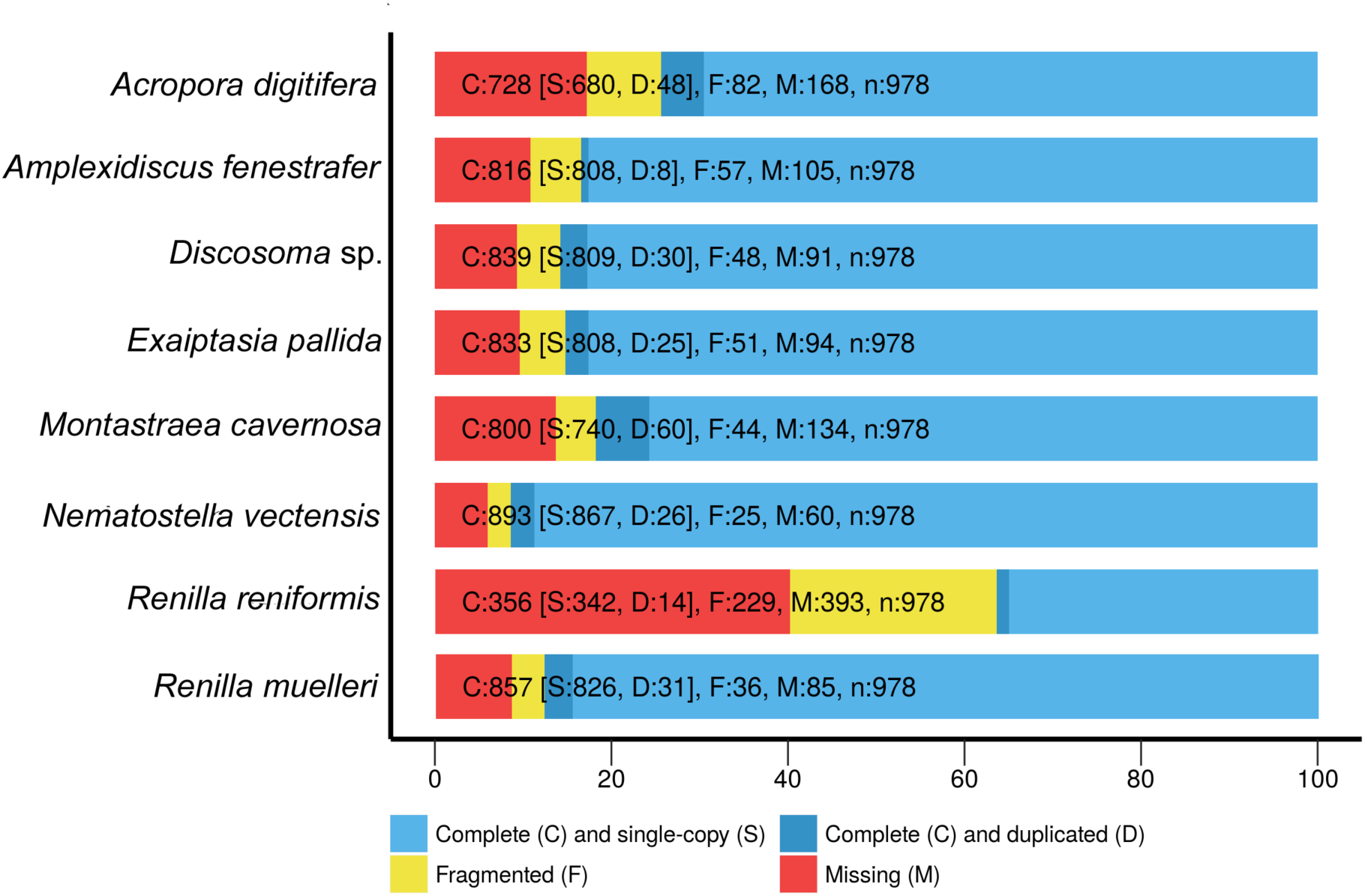
BUSCO-generated chart showing relative completeness of six hexacoral genomes, one octocoral genome, and the *Renilla muelleri* assembly.

**Supplemental Table S2.** *Renilla muelleri* genome assembly and annotation comparisons to other anthozoan genomes.

|  | Genome Size<br>(Mb) | Total #<br>Complete<br>BUSCOs** | Contig N50<br>(KB) | Exon length<br>(bp) | # Predicted<br>Gene models |
| --- | --- | --- | --- | --- | --- |
| *Acropora digitifera* | 420 | 728 | 10.6 | 230 | 23,668 |
| *Amplexidiscus fenestrafer* | 350 | 816 | 20.0 | 218 | 21,372 |
| *Discosoma* sp. | 428 | 839 | 18.7 | 226 | 23,199 |
| *Exaiptasia pallida* | 256 | 833 | 14.4 | NA | 26,042 |
| *Montastraea cavernosa* | 448 | 800 | 343 | NA | 30,360 |
| *Nematostella vectensis* * | 329 | 893 | 19.8 | 208 | 27,273 |
| *Renilla reniformis* | 132 | 356 | 1.8 | NA | 12,689 |
| *Renilla muelleri* | 172 | 857 | 64.8 | 249 | 23,360 |

\* Data taken from [52] and [56]
\*\*Complete BUSCOs generated from analysis herein

The number of predicted genes was highly similar across all anthozoan genomes (Supplementary Table S2). The range of predicted genes was 21,372 to 30,360 across the six hexacorals. The number of predicted genes (23,360) for *R. muelleri* was similar to the 23,668 genes predicted for *A. digitifera.*

Interestingly, the genome size of *R. muelleri* is considerably smaller (172 Mb) than other hexacoral genomes (256-448 Mb). Of the hexacorals, the anemone *Exaiptasia pallida* has the smallest genome size of 256 Mb, while the others have genome sizes >300 Mb. As indicated by [56], *E. pallida* has smaller and less frequent introns. Similar to *E. pallida,* exon sizes were larger in *R. muelleri* (249 bp) compared to the hexacorals (208 to 230 bp). These results suggest that there may be comparatively fewer non-coding regions in *R. muelleri* because the number of predicted gene models in *R. muelleri* is similar to hexacorals, yet the exon sizes are larger and the genome size is smaller in *R. muelleri.* In addition, repetitive elements in the *R. muelleri* genome may be less frequent, however, this remains to be further examined.

We also compared the mitochondrial genome to the previously published mitogenome of *R. muelleri* [59]. We used BLASTn to search for the mitogenome among the contigs (included as the last contig in the assembly) and recovered the entire 18,641 bp circularized, mitogenome. Compared to the published mitogenome, there were just two, single bp differences and one bp indel.

## Conclusions

We present the first octocoral genome assembly and showcase the feasibility of the MaSuRCA hybrid assembler for marine invertebrate genomics. The *R. muelleri* genome is one of the smallest anthozoan genomes discovered to date, yet it is comparable to other coral and anemone genomes in terms of predicted gene models. The identification of 88% of complete metazoan BUSCOs in the *R. muelleri* genome highlights that a quality genome assembly can be obtained from relatively low coverage sequencing of short and long read data. Although more data are needed to further increase size and reduce number of scaffolds, and further functional annotation is needed, the genome of the sea pansy, *R. muelleri,* provides a novel resource for the scientific community to further investigations of gene family evolution, comparative genomics, and the genomic basis of coral diversity.

## Availability of supporting data

The final hybrid assembly and predicted proteins generated by this study are in the *Giga*DB repository [60] and on the reefgenomics website [61]. Raw Illumina and PacBio reads are available in NCBI’s Sequence Read Archive (PRJNA491947). RNA-Seq reads have been uploaded to the European Nucleotide Archive (PRJEB28688).

## Abbreviations

bp: base pair
BUSCO: Benchmarking Universal Single-Copy Orthologs
Gb: gigabp
Mb: megabp
MY: million years
PE: paired end
Pacbio: Pacific Biosciences

## Additional Files

**Supplemental File 1.** Gene model annotations of *Renilla muelleri* using the *Nematostella vectensis* Joint Genome Institute filtered protein model.

**Supplemental File 2.** Gene annotations of *Renilla muelleri* using the *Nematostella vectensis* Vienna dataset.

**Supplemental File 3.** Reference file that includes annotations for the predicted gene models. Vienna dataset. This dataset includes GO terms, KOG IDs, and InterPro domains as annotated in the *Nematostella vectensis* filtered protein models (Joint Genome Institute).

## Authors’ Contributions

**Justin Jiang**: Conceptualization, Investigation, Formal Analysis, Software Programming, Methodology, Validation, Data Curation, Writing - Original Draft Preparation, Writing - Review & Editing, Visualization

**Andrea M. Quattrini**: Conceptualization, Supervision, Investigation, Formal Analysis, Methodology, Validation, Data Curation, Writing - Original Draft Preparation, Writing - Review & Editing, Visualization

**Warren R. Francis**: Software Programming, Methodology, Validation, Writing - Review & Editing

**Joseph F. Ryan**: Methodology, Validation, Data Curation, Writing - Review & Editing

**Estefania Rodriguez**: Conceptualization, Writing - Review & Editing

**Catherine S. McFadden**: Conceptualization, Supervision, Writing - Original Draft Preparation, Writing - Review & Editing

## Acknowledgments

We thank N. Alexander, C. Mason, and the Weill Cornell Medicine Epigenetics Core Facility and staff for MiSeq and PacBio sequencing. Thanks to M. Brugler, C. Schnitzler, and S. Herrera for advice. B. Macdonald generated the filterGenes.py script. We thank M. Heloski for collection of *Renilla reniformis* sample used for RNA-Seq.

## References

1. Daly M, Brugler MR, Cartwright P et. al. The phylum Cnidaria: A review of phylogenetic patterns and diversity 300 years after Linnaeus. Zootaxa. 2007;1668:127–182.

2. McFadden CS, France SC, Sánchez JA et. al. A molecular phylogenetic analysis of the Octocorallia (Cnidaria: Anthozoa) based on mitochondrial protein-coding sequences. Molecular Phylogenetics and Evolution. 2006;41(3):513:527.

3. Williams GC. The global diversity of sea pens (Cnidaria: Octocorallia: Pennatulacea). PLoS One. 2011;6:e22747

4. Williams GC. Living genera of sea pens (Coelenterata: Octocorallia: Pennatulacea): illustrated key and synopsis. Zoological Journal of the Linnean Society. 1995;113:93–140.

5. World Register of Marine Species: World List of Octocorallia Renillidae. http://marinespecies.org/aphia.php?p=taxdetails&id=266953, Accessed 19 Aug 2018.

6. Cairns SD, Bayer FM. Octocorallia (Cnidaria) of the Gulf of Mexico. In: Felder DL, Camp DK, editors. Gulf of Mexico–Origins, Waters, and Biota. Volume 1. Biodiversity. College Station, Texas: Academic; 2009:321–331.

7. Sherf BA, Navarro SL, Hannah RR, Wood KV. Dual-luciferase reporter assay: an advanced co-reporter technology integrating firefly and Renilla luciferase assays. Promega Notes. 1996;56:2.

8. Saito K, Chang YF, Horikawa K et al. Luminescent Proteins for High-Speed Single-Cell and Whole-Body Imaging. Nature Communications. 2012; doi:10.1038/ncomms2248.

9. Stepanenko OV, Verkhusha VV, Kuznetsova IM, Uversky VN, Turoverov KK. Current Protein & Peptide Science. 2008; doi:10.2174/138920308785132668

10. Clavico EE, De Souza AT, Da Gama BA, Pereira RC. Antipredator defense and phenotypic plasticity of sclerites from *Renilla muelleri*, a tropical sea pansy. The Biological Bulletin, 2007;213(2):135–140.

11. Ledoux JB, Antunes A. Beyond the beaten path: improving natural products bioprospecting using an eco-evolutionary framework–the case of the octocorals. Critical Reviews in Biotechnology. 2018;38(2):184–198.

12. Pop M, Salzberg SL. Bioinformatics challenges of new sequencing technology. Trends in Genetics. 2008;24(3):142–149.

13. Koren S, Schatz MC, Walenz BP, Martin J, Howard JT, Ganapathy G, Phillippy AM. Hybrid error correction and de novo assembly of single-molecule sequencing reads. Nature biotechnology, 2012;30(7):693.

14. English AC, Richards S, Han Y et al. Mind the gap: Upgrading genomes with Pacific Biosciences RS long-read sequencing technology. PLoS ONE. 2012; doi:10.1371/journal.pone.0047768.

15. Bashir A, Klamer AA, Robins WP et al. A hybrid approach for the automated finishing of bacterial genomes. Nature Biotechnology. 2012; doi:10.1038/nbt.2288.

16. Giordano F, Aigrain L, Quail MA et al. De novo yeast genome assemblies from MinION, PacBio and MiSeq platforms. Scientific Reports. 2017; doi:10.1038/s41598-017-03996-z.

17. Tan MH, Austin CM, Hammer MP et al. Finding Nemo: hybrid assembly with Oxford Nanopore and Illumina reads greatly improves the clownfish (*Amphiprion ocellaris*) genome assembly. GigaScience. 2018; doi:10.1093/gigascience/gix137.

18. McFadden CS, Alderslade P, Ofwegen LP van, Johnsen H, Rusmevichientong A. Phylogenetic relationships within the tropical soft coral genera *Sarcophyton* and *Lobophytum* (Anthozoa, Octocorallia). Invertebrate Biology 2006;125:288–305.

19. Bolger AM, Lohse M, Usadel B. Trimmomatic: a flexible trimmer for Illumina sequence data. Bioinformatics 2014;30(15):2114–20.

20. Wood DE, Salzberg SL. Kraken: ultrafast metagenomic sequence classification using exact alignments. Genome Biology. 2014;15(3):R46.

21. Wood DE. Minikraken 8 GB database, Johns Hopkins University, https://ccb.jhu.edu/software/kraken/dl/minikraken_20171019_8Gb.tgz (August 7 2018, date last accessed)

22. National Center for Biotechnology Information: Trivial HTTP: env_nt.00 to env_nt.23. ftp://ftp.ncbi.nlm.nih.gov/blast/db/

23. Boratyn GM, Camacho C, Cooper PS et al. BLAST: a more efficient report with usability improvements. Nucleic Acids Research 2013;41(W1):W29–33.

24. Huson DH, Mitra S, Ruscheweyh HJ et al. Integrative analysis of environmental sequences using MEGAN4, Genome Research, 2011;21:1552–1560.

25. Huson DH, Auch AF, Qi J et al. MEGAN analysis of metagenomic data, Genome Research, 2007;17(3):377–86.

26. Haas BJ, Papanicolaou A, Yassour M, Grabherr M, Blood PD, Bowden J, MacManes MD. De novo transcript sequence reconstruction from RNA-seq using the Trinity platform for reference generation and analysis. Nature protocol. 2013;8(8):1494.

27. Zimin AV, Marçais G, Puiu D et al. The MaSuRCA genome assembler. Bioinformatics 2013;29(21):2669–77.

28. Bankevich A, Nurk S, Antipov D, et al. SPAdes: A New Genome Assembly Algorithm and Its Applications to Single-Cell Sequencing; Journal of Computational Biology. 2012; doi:10.1089/cmb.2012.0021

29. Simão FA, Waterhouse RM, loannidis P et al. BUSCO: Assessing Genome Assembly and Annotation Completeness with Single-Copy Orthologs. Bioinformatics. 2015; doi:10.1093/bioinformatics/btv351.

30. Finn RD, Clements J, Eddy SR et al. HMMER Web Server: Interactive Sequence Similarity Searching. Nucleic Acids Research. 2011; doi:10.1093/nar/gkr367.

31. Bushnell B. BBMap Short Read Aligner. Berkeley, CA: University of California; 2016. https://sourceforge.net/projects/bbmap/ (August 7 2018, date last accessed).

32. Walker BJ, Abeel T, Shea T et al. Pilon: an integrated tool for comprehensive microbial variant detection and genome assembly improvement. PLoS One. 2014; doi:10.1371/journal.pone.0112963

33. Langmead B, Salzberg SL. Fast Gapped-Read Alignment with Bowtie 2. Nature Methods. 2012; doi:10.1038/nmeth.1923.

34. Francis WR haplotypeblastn.py; https://bitbucket.org/wrf/sequences/raw/f23b4dd3c965cc1774b9e10eb433242a18c13c65/haplotypeblastn.py (August 7 2018, date last accessed).

35. Hahn C select_contigs.pl; https://github.com/chrishah/phylog/blob/master/scripts-external/select_contigs.pl (August 7 2018, date last accessed).

36. Smit AFA, Hubley R, Green P. RepeatMasker; http://repeatmasker.org

37. Lunter G, Goodson M. Stampy: a statistical algorithm for sensitive and fast mapping of Illumina sequence reads. Genome Research. 2011;21(6):936–939.

38. Pena-Centeno T; filterBam, https://github.com/nextgenusfs/augustus/tree/master/auxprogs/filterBam

39. https://computationalbiologysite.wordpress.com/2013/07/25/incorporating-rnaseq-tophatto-augustus, (August 7 2018, date last accessed).

40. Stanke M, Steinkamp R, Waack S et al. AUGUSTUS: a web server for gene finding in eukaryotes. Nucleic Acids Research 2004;32(suppl–2):W309–12.

41. Stanke M, Schöffmann O, Morgenstern B, Waack S. Gene prediction in eukaryotes with a generalized hidden Markov model that uses hints from external sources. BMC Bioinformatics. 2006; doi:10.1186/1471-2105-7-62.

42. Francis WR, Wörheide G. Similar ratios of introns to intergenic sequence across animal genomes. Genome Biology and Evolution; 2017;9(6):1582–1598.

43. Joint Genomics Institute: Trivial HTTP, Nemve1. https://genome.jgi.doe.gov/portal/Nemve1/Nemve1.download.ftp.html (7 August 2018, date last accessed)

44. Macdonald B. filterGenes.py. https://github.com/mcfaddenlab/filterGenes.py/blob/master/README.md (August 7, 2018, date last accessed)

45. Uniprot Consortium. UniProt: the Universal Protein Knowledgebase. Nucleic Acids Research. 2018; doi:10.1093/nar/gky092

46. UniProt Consortium, Reviewed Swiss-Prot data, ftp://ftp.uniprot.org/pub/databp/uniprot/current_release/knowledgebase/complete/uniprot_sprot.fasta.gz (August 7, 2018, date last accessed)

47. https://ndownloader.figshare.com/files/1215191, (August 7 2018, date last accessed).

48. Moran Y, Fredman D, Praher D et al. Cnidarian MicroRNAs frequently regulate targets by cleavage. Genome Research. 2014; doi:10.1101/gr.162503.113.

49. Joint Genome Institute. Nematostella vectensis genome. Version 1. https://genome.jgi.doe.gov/portal/Nemve1/Nemve1.download.html (August 7, 2018, date last accessed).

50. Putnam NH, Srivastava M, Hellsten U, Dirks B, Chapman J, Salamov, A, et al. Sea anemone genome reveals ancestral eumetazoan gene repertoire and genomic organization. Science 2007;317(5834):86–94.

51. Shinzato C, Shoguchi E, Kawashima T et al. National Center for Biotechnology Information, Acropora digitifera genome Version 1. https://www.ncbi.nlm.nih.gov/nuccore/BACK00000000.1 (November 2015, date last accessed).

52. Shinzato C, Shoguchi E, Kawashima T, et al. Using the *Acropora digitifera* genome to understand coral responses to environmental change. Nature. 2011;476:7360–320.

53. Liew YJ, Aranda M, Voolstra CR. Reefgenomics.Org ‐ a repository for marine genomics data. Database (Oxford) 2016, 1–4 A*mplexidiscus fenestrafer* and *Discosoma* sp. genomes. http://corallimorpharia.reefgenomics.org (August 7, 2018, date last accessed).

54. Wang X, Liew YJ, Li Y, Zoccola D, Tambutte S, Aranda M. Draft genomes of the corallimorpharians *Amplexidiscus fenestrafer* and *Discosoma* sp. Molecular Ecology Resources 2017; 17(6); 187–195.

55. Baumgarten E, Simakov O, Esherick LY et al. National Center for Biotechnology Information, (*Ex*)*aiptasia pallida* genome Version 1.1 ftp://ftp.ncbi.nlm.nih.gov/sra/wgs_aux/LJ/WW/LJWW01/LJWW01.1.fsa_nt.gz (August 7 2018, date last accessed.

56. Baumgarten S, Simakov O, Esherick LY et al. The genome of *Aiptasia*, a sea anemone model for coral symbiosis. Proceedings of the National Academy of Sciences 2015;112(38):11893–11898.

57. Matz Lab. Montastraea cavernosa genome. Jul 2018 version. https://matzlab.weebly.com/data--code.html (August 7, 2018, date last accessed.

58. Kayal E, Bentlage B, Pankey MS et al. Phylogenomics provides a robust topology of the major cnidarian lineages and insights on the origins of key organismal traits. BMC Evolutionary Biology 2018;18:68.

59. Kayal E, Roure B, Phillipe H et al. Cnidarian phylogenetic relationships as revealed by mitogenomics. BMC Evolutionary Biology, 2013;13:5.

60. Jiang J, Quattrini AM, Francis WR, et al. A hybrid de novo assembly of the sea pansy (*Renilla muelleri)* genome. GigaScience Database 2018. doi:XXXXX

61. Liew YJ, Aranda M, Voolstra CR. Reefgenomics.Org ‐ a repository for marine genomics data. Database (Oxford) 2016, 1–4 *Renilla muelleri* genome http://rmue.reefgenomics.org (August 7, 2018, date last accessed)

